# The actin-capping protein alpha-adducin is required for T-cell costimulation

**DOI:** 10.1101/580191

**Authors:** Timothy J. Thauland, Manish J. Butte

**Affiliations:** Division of Immunology, Allergy, and Rheumatology, Department of Pediatrics, University of California, Los Angeles, CA, 90095 USA

## Abstract

Alpha-adducin (Add1) is a critical component of the actin-spectrin network in erythrocytes, acting to cap the fast-growing, barbed ends of actin filaments, and recruiting spectrin to these junctions. Add1 is highly expressed in T cells, but its role in T-cell activation has not been examined. Using a conditional knockout model, we show that Add1 is necessary for complete activation of CD4+ T cells in response to low levels of antigen but is dispensable for CD8+ T cell activation and response to infection. Surprisingly, costimulatory signals through CD28 were completely abrogated in the absence of Add1. This study is the first to examine the role of actin-capping in T cells, and it reveals a previously unappreciated role for the actin cytoskeleton in regulating costimulation.

## Introduction

Optimal T-cell activation requires the recognition of cognate peptide-MHC by the T-cell receptor (TCR) and additional signals from membrane-bound costimulatory receptors such as CD28. Ligation of CD28 by CD80 or CD86 results in recruitment of PI3K to its cytoplasmic tail and activation of the kinase Akt (Boomer and Green, 2010). Downstream targets of Akt include the kinase complex IKK, which activates the NF-κB pathway, leading to IL-2 production in T cells. CD28 is also required for positioning of PKC-θ at the immune synapse (Yokosuka et al., 2008). PKC-θ contributes to NF-κB activity via phosphorylation of CARD11 (Isakov and Altman, 2012), and assembly of the CARD11-Bcl10-MALT1 complex (Schulze-Luehrmann and Ghosh, 2006). Thus, multiple pathways downstream of CD28 promote NF-κB activation and IL-2 production in recently activated T cells.

T cells rapidly migrate through secondary lymphoid organs, and upon antigen recognition, reorient towards the antigen presenting cell (APC) and form an immune synapse. Both motility and immune synapse formation are actin-dependent processes, making the ability to rapidly reorganize the actin cytoskeleton a critical aspect of T-cell function. A number of cytoskeletal regulatory processes have been intensively studied in T cells, including WASP and Arp2/3-mediated branching and cofilin-mediated severing of F-actin (Burkhardt et al., 2008). Capping F-actin is another means of regulating the actin cytoskeleton. When actin capping proteins, which include CapZ (also known as capping protein) and the adducin family, bind to the fast-growing barbed end of F-actin, additional actin monomers are prevented from lengthening the filament. This activity grants cells another level of spatial control over actin polymerization.

Alpha-adducin (Add1) differs from CapZ in that it also functions to recruit spectrin to the actin cytoskeleton (Li et al., 1998). This activity links the dynamic actin cytoskeleton to the spectrin cytoskeleton, which underlies the plasma membrane. Among cells of the hematopoietic lineage, adducin proteins, which exist as heterodimers and tetramers of alpha-adducin with beta-or gamma-adducin, have been shown to be critical components of the cytoskeleton of platelets and red blood cells. In red blood cells, loss of Add1 causes a loss of structural integrity resulting in spherocytosis (Robledo et al., 2008). The role of Add1 in T-cell biology has not been examined despite its expression at high levels in lymphocytes.

The actin uncapping protein RLTPR (also known as CARMIL2) has been shown to play a crucial role in CD28-mediated T-cell costimulation (Liang et al., 2013). Interestingly, although RLTPR binds to and negatively regulates CapZ, this activity was recently shown to be dispensable for costimulation (Roncagalli et al., 2016). Thus, the role of actin-capping in T-cell activation, if any, is not clear. Here, we generated conditional knockout mice lacking Add1 in T cells. We show that CD4 T cells lacking Add1 develop normally but have a profound defect in CD28-mediated costimulation. This defect reduces T-cell proliferation in response to suboptimal concentrations of antigen and decreases cytokine production. These results introduce a new pathway by which T-cell costimulation is regulated by the cytoskeleton.

## Results and Discussion

T-cell activation has been shown to result in a decrease in Add1 after 24-72 hrs of stimulation, indicating that Add1 may play a preferential role in the biology of naïve T cells (Lu et al., 2004). We also observed a decrease in Add1 levels 24 hours after stimulation, however, CD4+ T-cell blasts expressed Add1 at levels similar to naïve T cells (**Fig. 1A**). These results indicate that while Add1 levels vary over the course of an immune response, the protein is present both in the naïve and effector states. Add1 is regulated by phosphorylation at multiple sites. Rho Kinase (ROCK)-mediated phosphorylation of Add1 at Thr445 and Thr480 enhances binding of Add1 to F-actin (Fukata et al., 1999; Kimura et al., 1998; Matsuoka et al., 2000), while F-actin-capping is inhibited by phosphorylation at its C-terminal MARCKS domain by members of the Protein Kinase C (PKC) family (Matsuoka et al., 1998, 2000). Given the importance of PKC-θ activity in T-cell function, we measured phosphorylation of Ser 724 in the MARCKS domain after TCR crosslinking. Add1 was rapidly phosphorylated upon T-cell activation (**Fig. 1B**). Phosphorylation peaked at 2-5 min and then declined to baseline by 30 min. Upon TCR triggering, T cells stop crawling and orient polymerization of the actin cytoskeleton towards the APC (Negulescu et al., 1996). These results suggest that a decrease in actin capping immediately facilitates this process.

**Figure 1.**
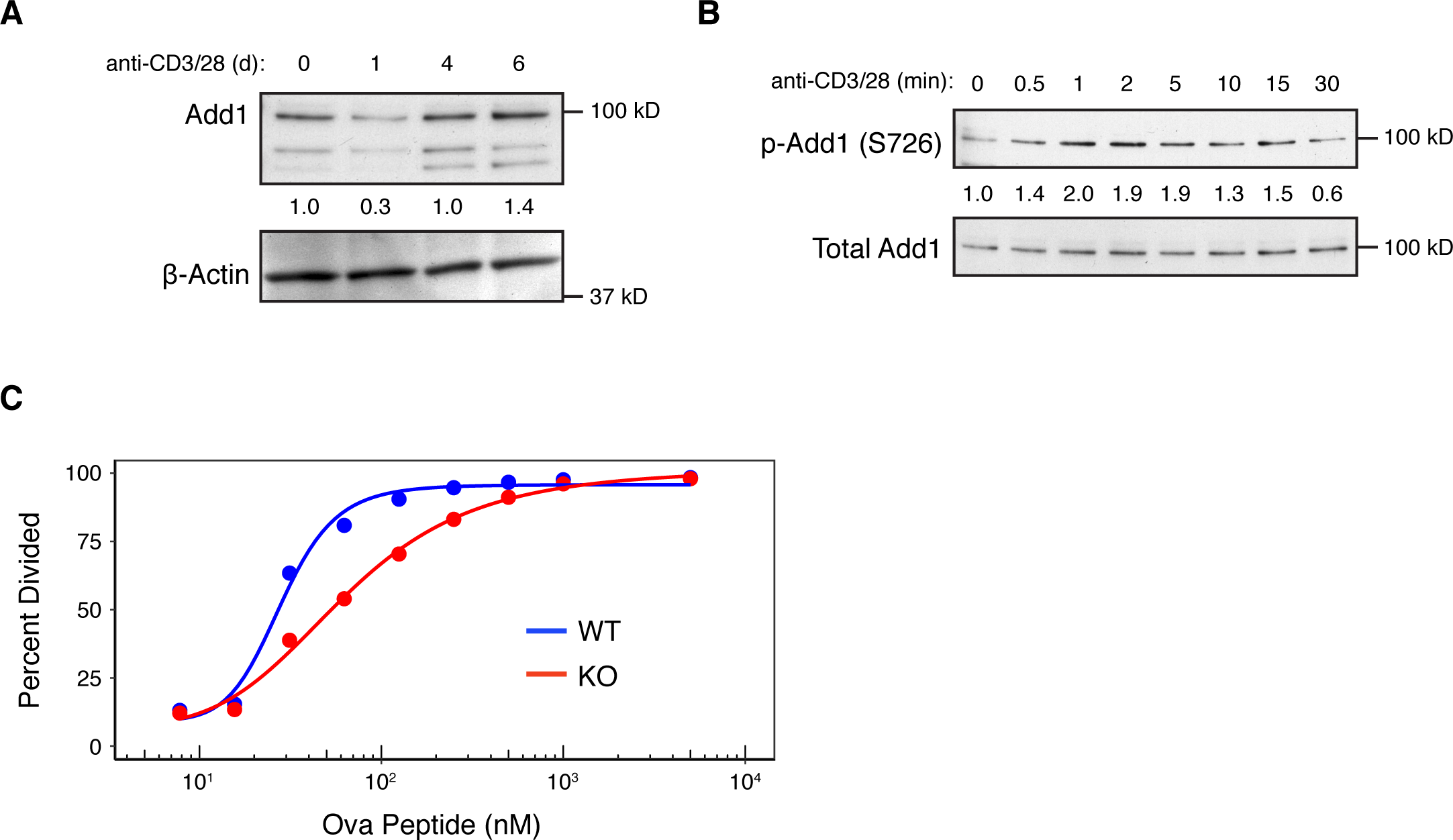
Add1 is phosphorylated upon TCR stimulation and is required for optimal CD4 T cell proliferation. **(A)** Expression of alpha-Adducin (Add1) was measured in CD4 T cells by western blot ex vivo and at d1, d4 and d6 after activation with anti-CD3 and anti-CD28. The two lower molecular weight bands are beta-and gamma-Adducin. **(B)** Phosphorylation of Add1 at Serine 726 was measured in naive CD4 T cells by western blot after anti-CD3 and anti-CD28 stimulation for the indicated times. **(C)** WT and Add1 KO OT-II CD4 T cells were labeled with CFSE and stimulated with LB27.4 B cells loaded with the indicated concentrations of Ova peptide. Cell division was measured on d3. Results are representative of two (A and B) or >5 experiments (C).

In order to examine the impact of Add1 on T-cell biology, we generated conditional knock-out (cKO) mice using the CD4-Cre system. T cells from these mice showed complete loss of alpha-adducin and also a loss of beta and gamma-adducin (**Fig. S1A**). This effect likely occurs because alpha-adducin pairs with either beta-or gamma-adducin to form stable heterodimers and has been previously noted in global Add1 KO mice (Robledo et al., 2008). Examination of T-cell development in the thymus revealed no defects in single positive, double positive, or double negative populations (**Fig. S1B and C**). Add1 KO mice also had normal percentages of CD4, CD8, CD44+CD62L-and FoxP3+ Treg cells in the periphery (**Fig. S1D-F**).

To study the role of Add1 in T-cell activation, we crossed Add1 cKO mice to TCR-transgenic OT-II mice that bear a TCR specific for Ova peptide presented by I-A^b^. When stimulated by APCs loaded with titrated doses of peptide, Add1 cKO cells exhibited defective proliferation as compared to controls (**Fig. 1C**). Interestingly, this effect was only seen at intermediate doses of peptide; at maximal TCR stimulation, Add1 KO CD4 T cells proliferated to the same degree as WT. This result suggested either a subtle defect in TCR signaling that only manifested at sub-optimal peptide doses, or a defect in costimulation.

To determine if Add1 plays a role in facilitating costimulation, we stimulated WT and KO CD4 T cells with various concentrations of anti-CD3 in the presence or absence of anti-CD28. At intermediate levels of TCR stimulation – where costimulation is most critical – Add1 KO T cells showed a profound impairment in CD28 costimulation as compared with WT. At very low or high concentrations of anti-CD3, WT and KO T cells proliferated equivalently, regardless of the presence of anti-CD28 (**Fig. 2A and 2B**). Measurement of CD25 upregulation at 48 hrs post-stimulation showed a similar effect; Add1 KO T cells showed defective upregulation of CD25 at moderate doses of anti-CD3 (**Fig. 2C and D**).

**Figure 2.**
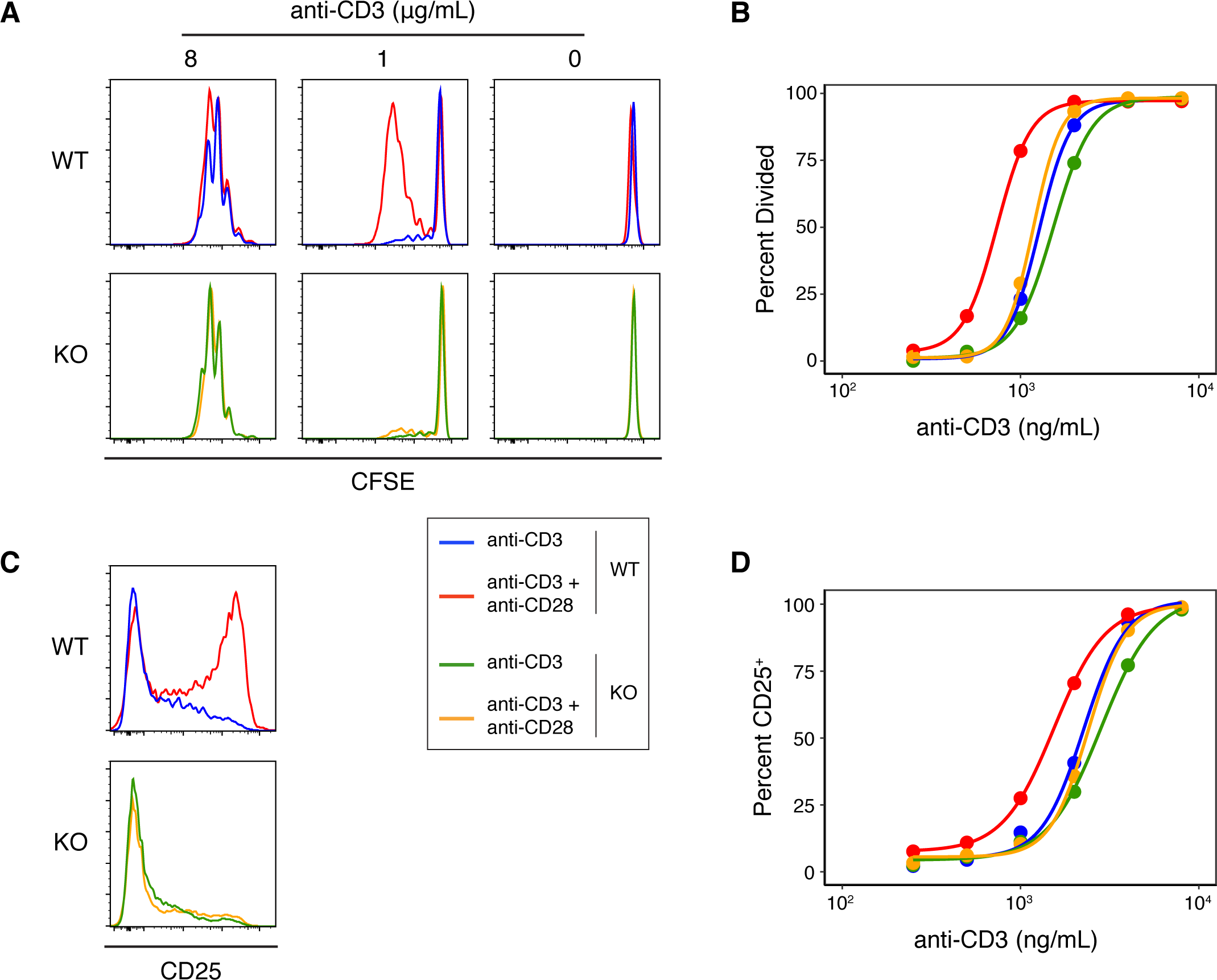
Add1 is required for CD28-mediated costimulation. **(A)** CFSE dilution of WT and KO CD4 T cells in response to the indicated concentrations of plate-bound anti-CD3 with or without soluble anti-CD28. Cell division was measured on d3. **(B)** Percent of cells that have divided at titrated doses of anti-CD3. **(C)** WT and Add1 KO OT-II CD4 T cells were stimulated with 2 μg/mL plate-bound anti-CD3 for 48 hr and assayed for CD25 expression. **(D)** Percent of cells positve for CD25 at titrated doses of anti-CD3. Results are representative of >5 (A and B) or two experiments (C and D).

Costimulatory signaling plays a critical role in IL-2 production (Greenwald et al., 2005), so we examined cytokine production by WT and KO CD4 T cells. We found that IL-2 production was impaired in KO cells, and this impairment was due to a failure to respond to costimulatory signals (**Fig. 3A**). KO cells also produced reduced amounts of IFN-γ as compared to WT, highlighting the importance of optimal stimulation for cytokine production (**Fig. 3B**).

**Figure 3.**
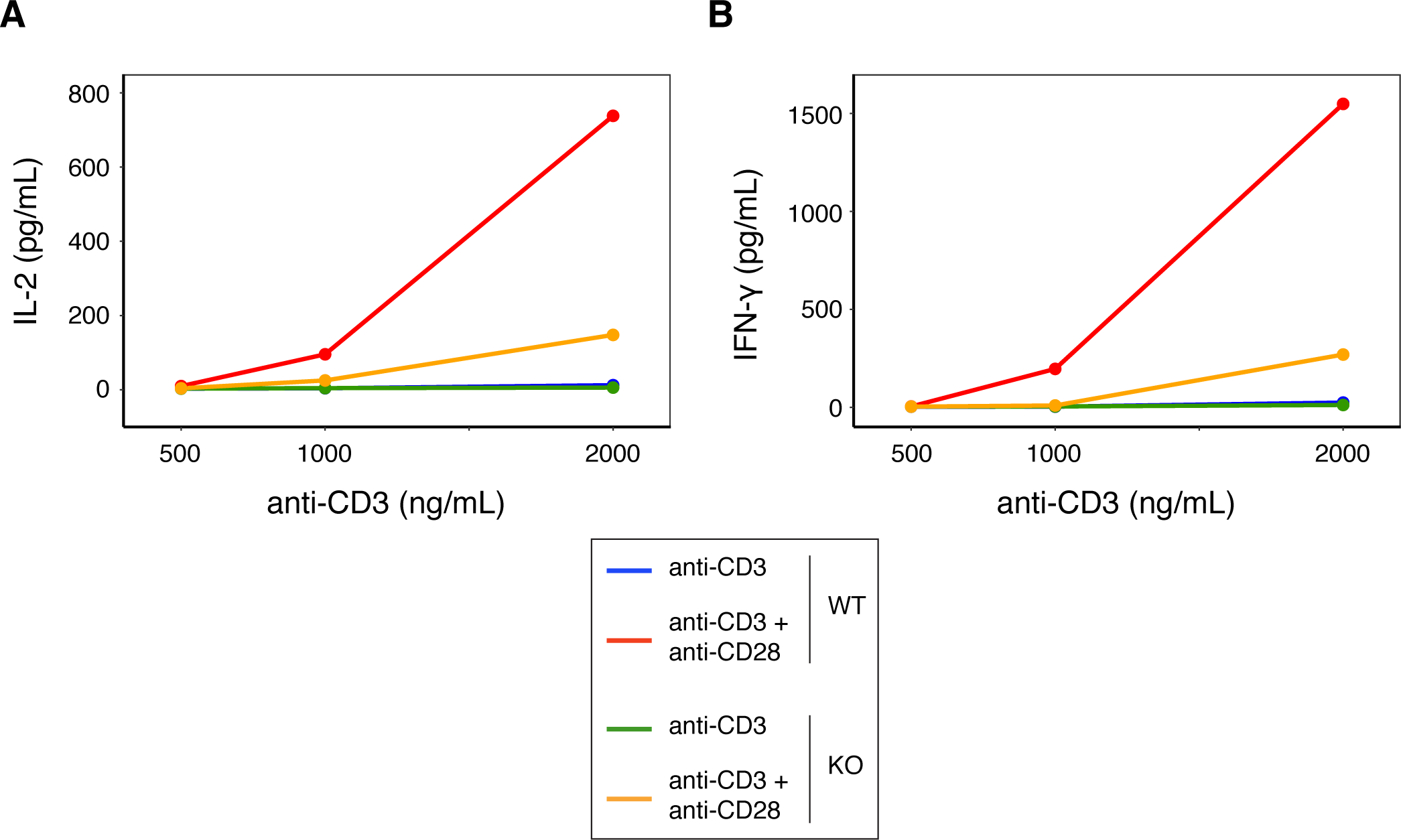
Add1 is required for IL-2 and IFN-γ production. **(A)** IL-2 and **(B)** IFN-γ production by Add1 WT and KO CD4 T cells stimulated with titrated amounts of anti-CD3 with and without soluble anti-CD28. Cytokine concentrations were measured from supernatants on day 2 of activation by Cytokine Bead Array. Results are representative of two experiments.

We tested whether a similar proliferative defect existed in CD8 T cells by stimulating with a range of anti-CD3 concentrations, but we saw no difference between the proliferative capacity of WT and KO T cells (**Fig. 4A**). To test the role of Add1 in CD8 cells *in vivo*, we crossed Add1 KO mice with OT-I TCR transgenic mice. We simultaneously infected CD45.1^+^ congenic mice with *Listeria monocytogenes* engineered to express Ova peptide and transferred naïve WT or Add1 KO CD45.2^+^ OT-I cells. Eight days post-infection, we harvested spleens from recipient mice and measured the accumulation and differentiation of donor T cells. We saw no difference in accumulation (**Fig. 4B and C**) or differentiation to KLRG1+CD127-short-lived effectors (**Fig. 4D and E**) and modestly reduced differentiation to KLRG1-CD127+ memory-precursor cells (**Fig. 4D and F**). We also measured cytokine production *ex vivo* and found that similar percentages of WT and KO cells made IFN-γ (**Fig. 4G and H**) and slightly fewer KO cells made TNF-α (**Fig. 4G and I**). These results could be explained by the fact that naïve CD8 T cells are not as dependent on costimulation as CD4 T cells (Wang et al., 2014). Given these results, it is likely that the role of Add1 in T-cell activation is primarily related to costimulation and not TCR triggering.

**Figure 4.**
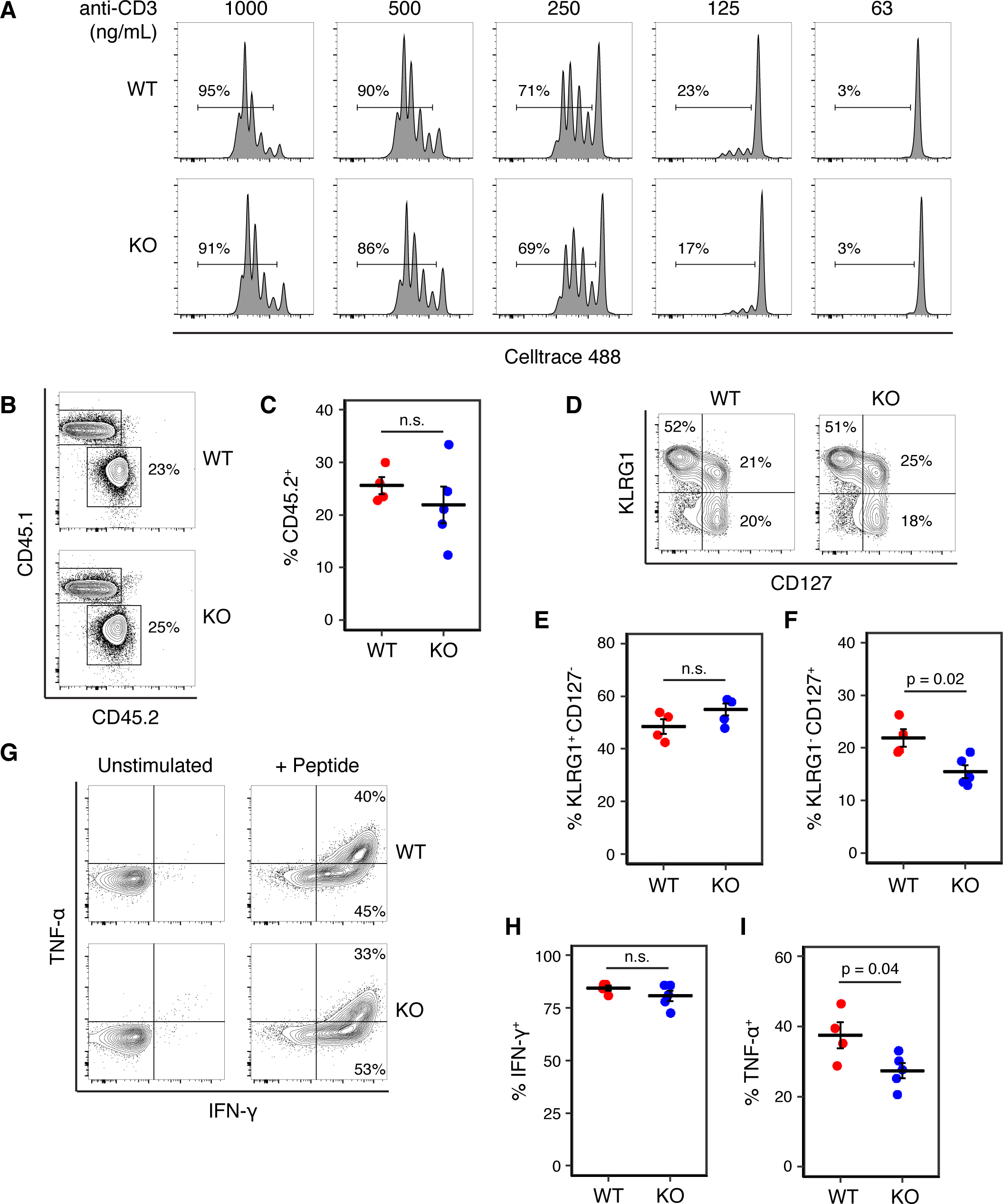
Add1 is not required for CD8 T cell activation or response to Listeria infection. **(A)** CD8 T cells from WT and Add1 cKO mice were stimulated with the indicated concentrations of anti-CD3. **(B and C)** B6 CD45.1 recipient mice were infected with 3,000 CFU of Lm-Ova and 10^5^ OT-I CD8 T cells from WT or KO mice. On d8, accumulation of donor CD45.2+ cells was measured (B) and quantified for multiple recipients (C)**. (D-F)** Among transferred cells, the percentage with a KLRG1+CD127-short-lived effector phenotype and KLRG1-CD127+ memory-precursor phenotype were measured (D) and quantified (E and F). **(G-I)** Splenocytes from infected mice were stimulated with Ova peptide and intracellular cytokine staining for IFN-γ and TNF-α was performed. The percentage of IFN-γ+ and TNF-α+ donor cells was measuredby FACS (G) and quantified (H and I). Results are representative of two experiments.

To our knowledge, this the first study identifying a role for the actin cytoskeleton in facilitating CD28-mediated costimulation. The actin-uncapping protein RLTPR is critical for costimulation, but only the scaffolding function was required for this role (Roncagalli et al., 2016). On the other hand, there is well established role for costimulation in regulating F-actin dynamics at the immune synapse. Blocking CD28 signaling affects the accumulation and centralization of TCR-pMHC clusters and optimal F-actin accumulation at the immune synapse (Tskvitaria-Fuller et al., 2014). CD28 costimulation appears to impact F-actin dynamics in multiple ways. WAVE2 and HS1, which control actin nucleation, and cofilin, which severs F-actin, showed defective accumulation at the synapse upon costimulation blockade (Roybal et al., 2016). Further, the actin nucleation promoting protein WASp has been shown to promote CD28-mediated F-actin accumulation and CD28 endocytosis (Badour et al., 2007). CD28 associates with filamin A, the actin-bundling protein and scaffold for lipid-raft nucleation, at the immune synapse, leading to recruitment of PKC-θ (Tavano et al., 2006; Hayashi and Altman, 2006) In the context of these reports, our results indicate that CD28 signaling and F-actin dynamics reciprocally regulate one another.

The mechanism by which Add1 facilitates costimulation will be an important area of future study. It is possible that the actin-capping properties of Add1 are required to correctly orient CD28 at the immune synapse. CD28 forms an annular ring around TCR-pMHC microclusters at the immune synapse and lead to recruitment of PKC-θ (Yokosuka et al., 2008). We hypothesize that actin-capping is a requirement for this process. Given that Add1 links the actin and spectrin cytoskeletons, it is tempting to speculate that reorganization of spectrin plays a role in costimulation. Indeed, antigen stimulation induces membrane-localization of α-spectrin in CD4 T cells (Lee et al., 1988), and α-II-spectrin has been shown to rapidly reorient to the immune synapse where it colocalizes with F-actin (Meissner et al., 2017). Further experiments will be necessary to dissect the relative importance of Add1-mediated actin-capping and spectrin recruitment in costimulation.

## Acknowledgements

We gratefully acknowledge support from the NIH (R01 gm110482).

## Materials and Methods

### Mice

C57BL/6 (Strain 000664), CD45.1 (Strain 002014), OT-II TCR transgenic (Strain 004194) and OT-I TCR transgenic mice (Strain 003831) were acquired from Jackson Labs. We acquired C57BL/6N embryonic stem (ES) cells that had been injected with an Add1 knockout first allele containing a neomycin selection cassette surrounded by FRT sites and upstream of LoxP sites surrounding exon 2 of Add1 from the European Conditional Mouse Mutagenesis program, as part of the Knock Out Mouse Project. ES were expanded and implanted into B6 albino mice and the resulting pups were eventually bred to C57BL/6 mice. Progeny were genotyped to determine if the transgene was transmitted and were bred to FLP deleter mice to remove the selection cassette (JAX Strain 009086). Add1 conditional KO mice were then generated by breeding to CD4-Cre (JAX Strain 022071).

### Cell lines and reagents

The I-A^b^+ B cell lymphoma line LB27.4 was purchased from the American Type Culture Collection. The following antibodies were used in this study: Fc Block, CD8a BV 421, CD45.2 BV 510, B220 BV 605, CD127 AF 488, CD45.1 PE, KLRG1 PE-Cy7, CD25 AF 647, TNF-α AF 488 and IFN-γ AF 647 were from Biolegend. CellTrace 488, CellTrace Violet and CellTrace CFSE were from ThermoFisher. Alpha-adducin antibody was from Abcam (ab40760). Phospho-alpha-adducin (Ser 726) antibody was from Santa Cruz Biotechnology. Class I (257-264) and Class II (323-339) ovalbumin peptides were from AnaSpec. GolgiPlug was from BD Biosciences. Anti-CD3 (145-2C11) and anti-CD28 (37.51) were from Bio-X-Cell.

### Western Blots

CD4 T cells were lysed in RIPA buffer containing Halt Protease inhibitor cocktail (Life Technologies). Soluble fractions of lysates were run on SDS–polyacrylamide gels and transferred to PVDF membranes. Membranes were blocked with 5% bovine serum albumin and incubated with the appropriate primary and HRP-conjugated secondary antibodies (GE Healthcare). Western Lightning ECL Pro (PerkinElmer) was used as the chemiluminescent substrate. For the acute stimulation experiment, naïve CD4 T cells were coated with 10 μg/mL anti-CD3 on ice for 10 min, washed and incubated at 37 °C for various times in the presence of 50 μg/mL goat anti-hamster crosslinking antibody. Cells were quickly centrifuged and lysed as above in the presence of PhosphataseArrest I (G-Biosciences).

### T-cell proliferation assays

CD4 or CD8 T cells were purified from the spleens of Add1 cKO mice using EasySep negative selection (StemCell). Cells were loaded with Celltrace 488 or Celltrace Violet and grown on 96 well plates pre-coated with the indicated concentrations of anti-CD3, with or without anti-CD28. Alternatively, OT-II^+^ CD4 T cells were loaded with Celltrace and incubated with LB27.4 B cells loaded with the indicated concentrations of Ova peptide. Cells were assayed by flow cytometry on day 3.

### Cytokine measurements

CD4 T cells were stimulated with plate-bound anti-CD3, with or without anti-CD28, as above. At 48 hr, supernatants were collected, and IL-2 and IFN-γ concentrations were measured with Cytokine Bead Array reagents (BD Biosciences), following the manufacturer’s instructions.

### Listeria Infections

OT-I^+^ CD8 T cells were purified from the spleens of WT or Add1 KO mice using EasySep negative selection. 10^5^ T cells and 3,000 colony forming units of *Listeria monocytogenes* expressing ovalbumin (a kind gift from Max Krummel). T cells and bacteria were injected *i.v*. into CD45.1^+^ recipient mice. On d8 post-infection, spleens were harvested from recipient mice and red blood cells were lysed. Some splenocytes were immediately surface stained for CD8a, B220, CD45.1, CD45.2, CD127, and KLRG1 and analyzed by flow cytometry to measure accumulation of donor CD45.2+ cells and development of short-lived effector and memory-precursor cells. To measure cytokine production, 10^7^ splenocytes were cultured with 10 μM Ova peptide for 6 hrs in the presence of GolgiPlug for the final 3 hrs. Cells were surface stained for CD8a, B220, CD45.1 and CD45.2 and then fixed with 2% paraformaldehyde. Cells were permeabilized with 0.5% saponin, stained for TNF-α and IFN-γ, and analyzed by flow cytometry.

