## Supplemental Figure 1 for "The actin-capping protein alpha-adducin is required for T-cell costimulation"

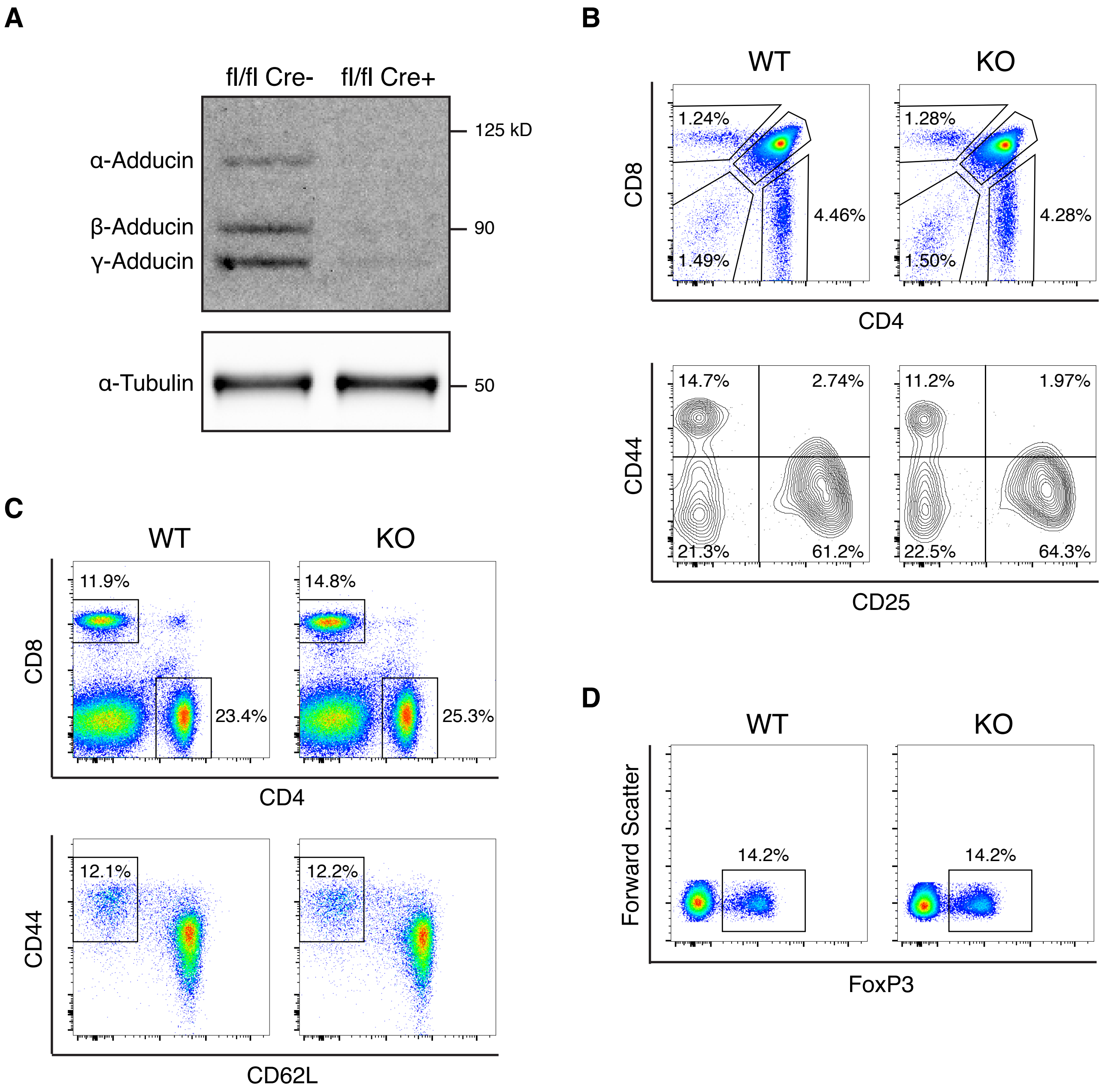


**Figure S1. T cell development is normal in the absence of Add1. (A)** Expression of alpha-Adducin (Add1) was measured in WT and KO CD4 T cells purified from the spleen. **(B)** Single positive and double positive (Top) and double negative (Bottom) populations in the thymus of WT and cKO mice were measured by FACS. **(C)** CD4 and CD8 populations and CD4+CD44+CD62L- memory cells were measured in the spleens of WT and Add1 cKO mice. **(D)** Percentage of FoxP3+ cells among splenic CD4+ cells from WT and Add1 cKO mice were measured.
